# APOBEC5: characterisation of a novel member of the AID/APOBEC protein family

**DOI:** 10.1101/2025.05.21.655229

**Authors:** Shuoshuo Tian, Paula Ellenberg, Betty Kouskousis, Gilda Tachedjian, Joshua A Hayward

## Abstract

The AID/APOBEC protein family is involved in diverse biological processes, most notably antiviral innate immunity, a function carried out by APOBEC3 (A3) in placental mammals. A3 is exclusive to this lineage, having emerged after the divergence from marsupials. This raises the question of how marsupials, which lack A3, defend against retroviruses. An uncharacterized A3 homologue, APOBEC5 (A5), is present in marsupial genomes and in some other vertebrate taxa. Here, we use in silico and in vitro approaches to investigate whether A5 serves as a functional antiviral counterpart to A3 in marsupials and whether marsupial genomes contain evidence of past A3-like activity against retroviral infections. We find that A5 was present in the last common ancestor of all jawed vertebrates but has been lost independently in multiple lineages. A5 exhibits unique structural and post-translational features not observed in APOBEC3 or other APOBEC proteins and has a broad subcellular and tissue distribution, suggesting a multifunctional role. Mutagenesis assays demonstrate that A5 functions as a DNA mutator and modestly restricts the infectivity of a model retrovirus, HIV-1, and that this activity is counteracted by the HIV-1 protein, Vif. Furthermore, analysis of the gray short-tailed opossum genome reveals distinct patterns of A3-like restriction in two groups of recently integrated retroviruses, providing direct evidence of an ongoing evolutionary conflict between marsupial APOBEC proteins and retroviruses. This study presents the first characterisation of A5 as a novel AID/APOBEC family member and reveals that marsupials possess an antiviral function homologous to placental A3, shedding light on the evolutionary dynamics of vertebrate antiviral immunity.

## Introduction

APOBEC3 (A3; apolipoprotein B mRNA editing enzyme, catalytic subunit 3) proteins are exclusive to placental mammals and play a crucial role as members of the antiviral innate immune response (Münk, et al. 2012; Stavrou and Ross 2015). A3 proteins are members of the multifunctional AID/APOBEC protein family of zinc-dependent cytosine deaminases which play diverse roles in processes including metabolism, immunity, cancer development, and viral defence (Harris and Dudley 2015; Mertz, et al. 2022; Pecori, et al. 2022). A3 proteins exert their antiviral effect by interfering with viral nucleic acid strand synthesis, primarily by the catalytic conversion of cytosine (C) to uracil (U) in single-stranded DNA (Chiu and Greene 2008). The importance of A3 proteins in the mammalian antiviral response is emphasised by the expansion and diversification of *A3* genes in many animals, including humans, non-human primates, and bats (Hayward, et al. 2018). A3 proteins impact the replication of diverse viruses but are best characterised for their role in targeting the replication of retroviruses, including the human immunodeficiency virus (HIV) (Uriu, et al. 2021).

Retroviruses infect all animal taxa, though most well characterised retroviruses are those that infect mammals. The evolution and diversification of A3 to become a critical antiretroviral defence in placental mammals begs the question of how the other extant mammalian lineages, marsupials and monotremes, mount their defence against retroviruses in the absence of an A3 repertoire. All mammals possess APOBEC1 (A1), APOBEC2 (A2), APOBEC4 (A4) and AID (activation-induced deaminase) (Salter, et al. 2016). Each of these have primary roles unrelated to antiviral immunity. A1 edits mRNA and plays a role in lipid metabolism (Salter, et al. 2016), A2 is involved in muscle development and maintenance (Sato, et al. 2010), A4 is hypothesized to have a role in spermatogenesis and regulation of gene expression (Marino, et al. 2016; Shi, et al. 2020), and AID is involved in antibody diversification (Conticello, et al. 2007).

Previous studies have noted that the genomes of marsupial mammals contain a gene that appears to be homologous to *A3*, referred to as *APOBEC5* (*A5*) or uncharacterised APOBEC (*UA2*) (Severi, et al. 2011; Ito, et al. 2020). A5 is absent from placental mammals, and is speculated to serve a similar antiviral function as A3 (Severi, et al. 2011; Ito, et al. 2020). *A5* genes were also noted to be present beyond mammals, in the genomes of a frog and the anole lizard (Severi, et al. 2011). Apart from these initial reports, A5 remains uncharacterised, and its function unknown. In this study, we characterised marsupial A5 to address the hypothesis that it represents a functional antiviral homologue of placental A3. We also analysed endogenous retroviruses (ERVs), within the genome of the gray short-tailed opossum, to determine if retroviruses that originally infected marsupials have been subjected to A3-like restriction.

## Results

### A5 is a unique member of the APOBEC family lost from several vertebrate lineages and possesses key conserved sequence and structural motifs

To reassess claims that A5 is a novel member of the APOBEC family and assess its presence in various animal taxa, we searched for publicly accessible APOBEC-like sequences in GenBank by BLAST homology to the previously reported A5 from the Tasmanian devil. This search revealed sequences homologous to previously reported ‘A5/UA2’ proteins in marsupials, reptiles, and amphibians, but not in any monotreme or placental mammal, consistent with previous reports (Severi, et al. 2011; Ito, et al. 2020). Additionally, this search revealed A5 homologues for the first time in numerous species of jawed fish, but not birds, which diverged later. Three species of sturgeon were also found to possess a duplication of A5 (referred to here as A5A and A5B) (Figure 1A-B).

**Figure 1.**
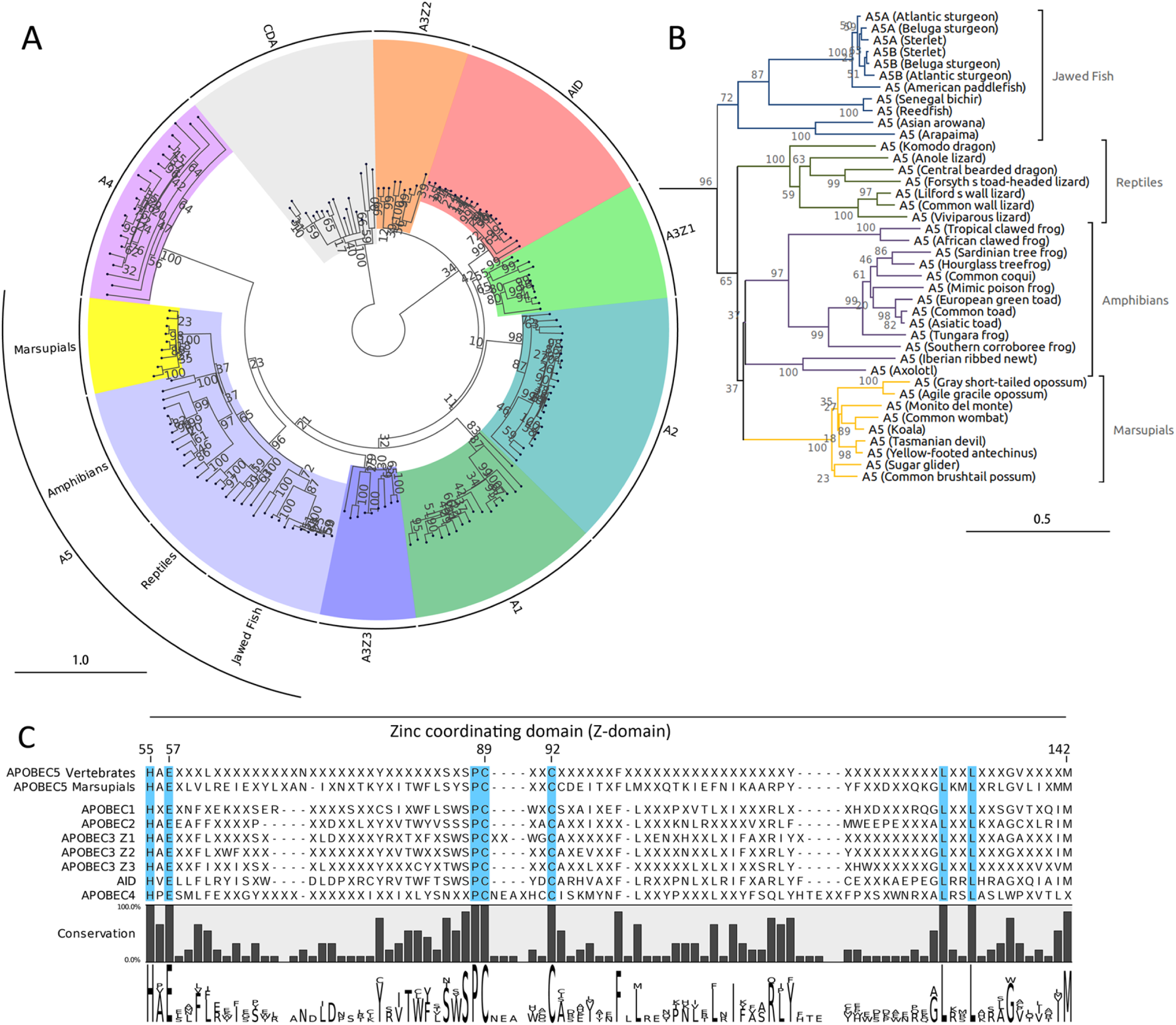
APOBEC5 (A5) phylogeny and sequence analysis.A. The phylogenetic relationships of candidate A5 amino acid sequences and taxonomically diverse representatives of each CDA and AID/APOBEC clade were inferred using the maximum likelihood method and JTT+G+T model with 1,000 bootstrap replicates. **B** The A5 subtree representing 36 jawed vertebrate species is shown. Scales in A and B represent the number of amino acid substitutions per site. **C** The consensus sequences of the zinc-coordinating domain (Z-domain) of each AID/APOBEC clade are shown. Conserved amino acid residues represented in >80% of analysed sequences are represented by their letter. ‘X’ denotes sites not represented by the same amino acid in >80% of species. Sites conserved in 100% of analysed sequences are highlighted in blue. The amino acid logo displays the various residues represented at each site with a height proportional to the percentage of their conservation. Amino acids are numbered relative to their position in Tasmanian devil A5. CDA, cytidine deaminase; AID, activation-induced deaminase.

Phylogenetic analysis of these sequences alongside other AID/APOBEC proteins and more distantly related cytidine deaminase (CDA) proteins confirmed their position among the APOBEC family. A5 was the shortest protein sequence distance from the A4 clade (Figure 1A, Supplementary table 1). A5 sequences from marsupials, reptiles, amphibians, and jawed fish formed discrete clades consistent with the divergence of host taxa (Figure 1B).

Analysis of the putative zinc-coordinating domain (Z-domain) of A5 sequences revealed that all A5 sequences contained the canonical HxEx_y_PCx_2-5_C Z-domain core sequence motif of AID/APOBEC proteins (Salter, et al. 2016; Ito, et al. 2020) (Figure 1C), suggesting conservation of the nucleic acid binding and/or cytosine deamination function. The conserved A5 Z-domain was revealed to be HAEx_3_Lx_24_SxSPCxxCx_6_Fx_17_Yx_10_LxxLx_3_GVx_4_M with >80% conservation across 36 diverse vertebrate species across all taxa possessing A5 (Figure 1C).

As AID/APOBEC proteins share similar structures (Salter, et al. 2016), we assessed whether A5 proteins were predicted to be structurally similar to other AID/APOBECs and potentially capable of physically interacting with nucleic acids. Using AlphaFold 3 (Abramson, et al. 2024), the structure of A5 proteins representing marsupials, reptiles, amphibians, and jawed fish were predicted in silico (Figure 2, marsupial Tasmanian devil A5; Additional A5 structures, Supplementary Figure 1). This analysis predicted A5 forming an A5-zinc-ssDNA complex, including a U-shaped DNA substrate conformation, with strong similarity to human A3 proteins (Salter, et al. 2016; Kouno, et al. 2017) (Figure 2A).

**Figure 2.**
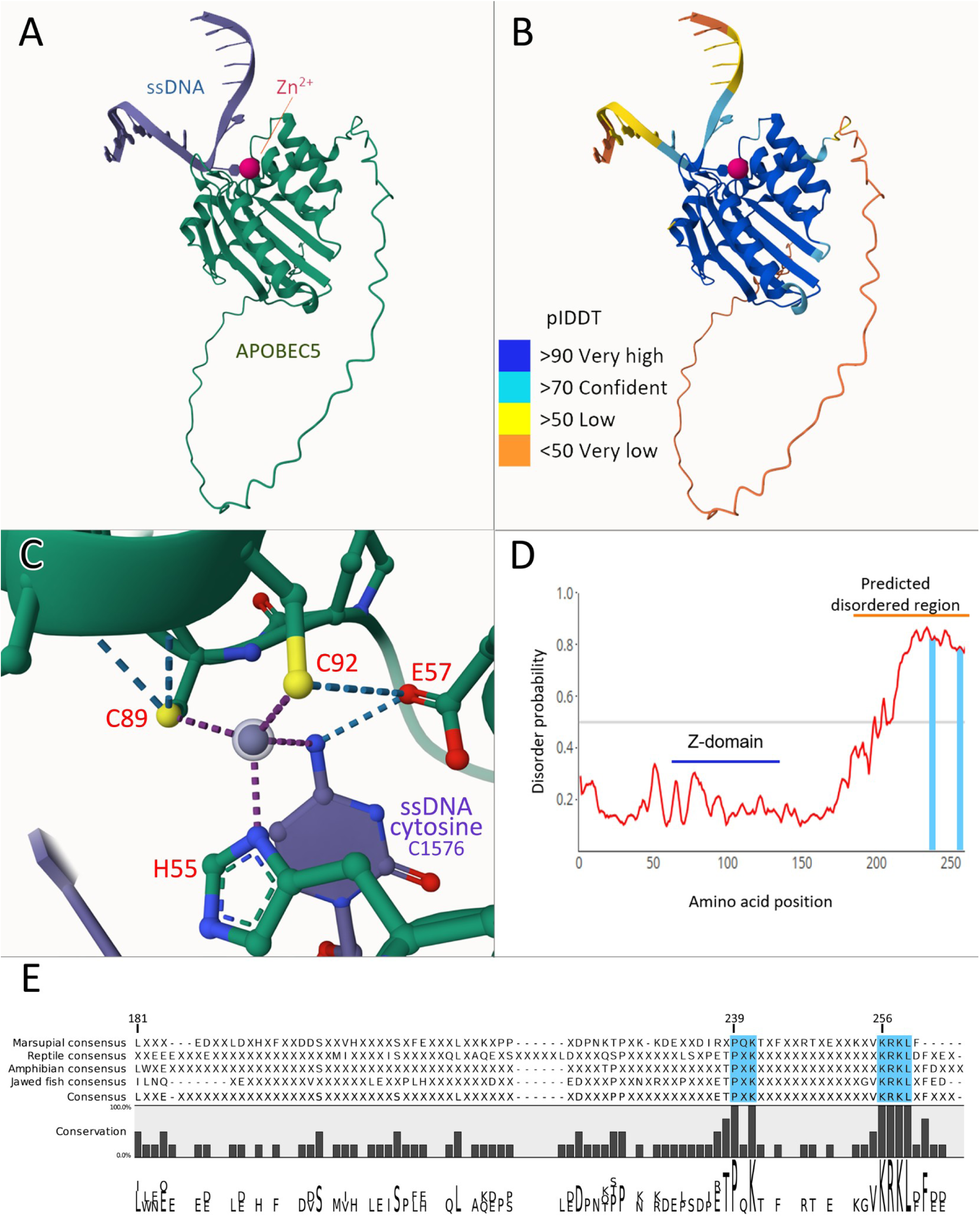
Structural prediction of APOBEC5 (A5) with Alpha Fold 3 and AIUPred. **A** Prediction of A5 structure of Tasmanian devil A5 (tdA5, green), complexed with a zinc ion (pink) and a 15-mer ssDNA sequence (5’-GATTACGCACAAACG-3’, blue). The ssDNA and zinc ion form a complex with A5 at the zinc-coordinating domain (Z-domain). **B** A confidence-scored representation of the tdA5-Zn-sDNA structure, with residues coloured according to the confidence of the predicted local structure. The Alpha Fold 3 predicted Local Distance Difference Test (pLDDT) score is the measure of confidence, and values are indicated in the key **(inset)**. **C** Close-up view of the Z-domain with the bound amine group of a cytosine within the ssDNA. Conserved histidine (H), cysteine (C), and glutamic acid (E) residues (in red text) and the bound cytosine (ssDNA cytosine c1576) are shown. **D** A representative AIUPred prediction of disorder (probability > 0.5) at each amino acid position for tdA5. **E** Multiple sequence alignment of the consensus sequences for the C-terminal predicted disordered region of A5 for marsupials, reptiles, amphibians, and jawed fish reveal conserved PxK and KRKL motifs (highlighted in blue). Conserved amino acid residues represented in >80% of A5 sequences are represented by their letter. ‘X’ denotes sites not represented by the same amino acid in >80% of A5 sequences. Amino acids are numbered relative to their position in tdA5.

The N-terminal region of A5, which contains the Z-domain, was predicted to form a canonical cytidine deaminase (CDA) fold structure (Salter, et al. 2016) with very high confidence, while the C-terminal region was not predicted to possess an intrinsically stable 3D structure (Figure 2B). The A5 protein, zinc ion, and ssDNA were predicted to form a complex at the Z-domain (Figure 2C), with the core H55, E57, C89, and C92 conserved residues forming metal coordination bonds directly with the zinc ion and hydrogen bonds with each other, and the glutamic acid (E57) residue hydrogen bonding directly to the amine group of a cytosine within the ssDNA molecule. These data indicate that A5 is likely a structural homologue of A3 proteins, as these features are observed for A3 proteins (Salter, et al. 2016; Kouno, et al. 2017) (Supplementary Figures 1 & 2). Additionally, Alpha Fold 3 predictions of AID/APOBEC1-4 proteins of placental and marsupial mammals bear strong similarity to each other and to A5 (Supplementary Figures 1 & 2). Across different taxa, A5 structure predictions were highly similar to each other, with all structures predicted to possess a CDA structure with high confidence and an unstructured C-terminal region (Supplementary Figure 1).

To support the prediction of a lack of stable structure in the A5 C-terminal region, we further analysed A5 sequences using AIUPred (Erdős and Dosztányi 2024) to determine whether A5 was likely to contain an intrinsically disordered domain. In agreement with Alpha Fold 3, AIUPred calculated that A5 proteins contained an intrinsically disordered region at the C-terminal (Figure 2D, Supplementary Figure 3). AIUPred analysis of other APOBEC proteins revealed that A4 similarly possesses an extended, intrinsically disordered C-terminal region (Supplementary Figure 3). Sequence analysis of the predicted disordered region of A5 across each vertebrate taxa indicated a low degree of sequence conservation with the exception of two highly conserved protein motifs, 239PxK and 256KRKL at the far end of the unstructured C-terminal region (Figure 2E), suggesting that these motifs provide a conserved function.

Taken together, sequence, phylogenetic, and predictive structural analyses strongly suggest that A5 is a bona fide APOBEC protein and functional cytosine deaminase, that was present in the last common ancestor of jawed vertebrates which has been lost from multiple extant lineages, including placental mammals.

### Marsupial A5 is highly expressed in the colon and localises to both the cytoplasm and nucleus, suggesting a potential multi-functional role

To confirm that the marsupial *A5* gene is actively expressed in vivo, we assessed the relative tissue expression in the opossum by measuring A5 mRNA read abundance across different tissues within a publicly accessible gray short-tailed opossum transcriptome from multiple individuals (GenBank BioProject: PRJNA200320). Opossum A5 was broadly expressed across all analysed tissues, but most highly expressed in the colon, followed by the kidneys and liver (Figure 3).

**Figure 3.**
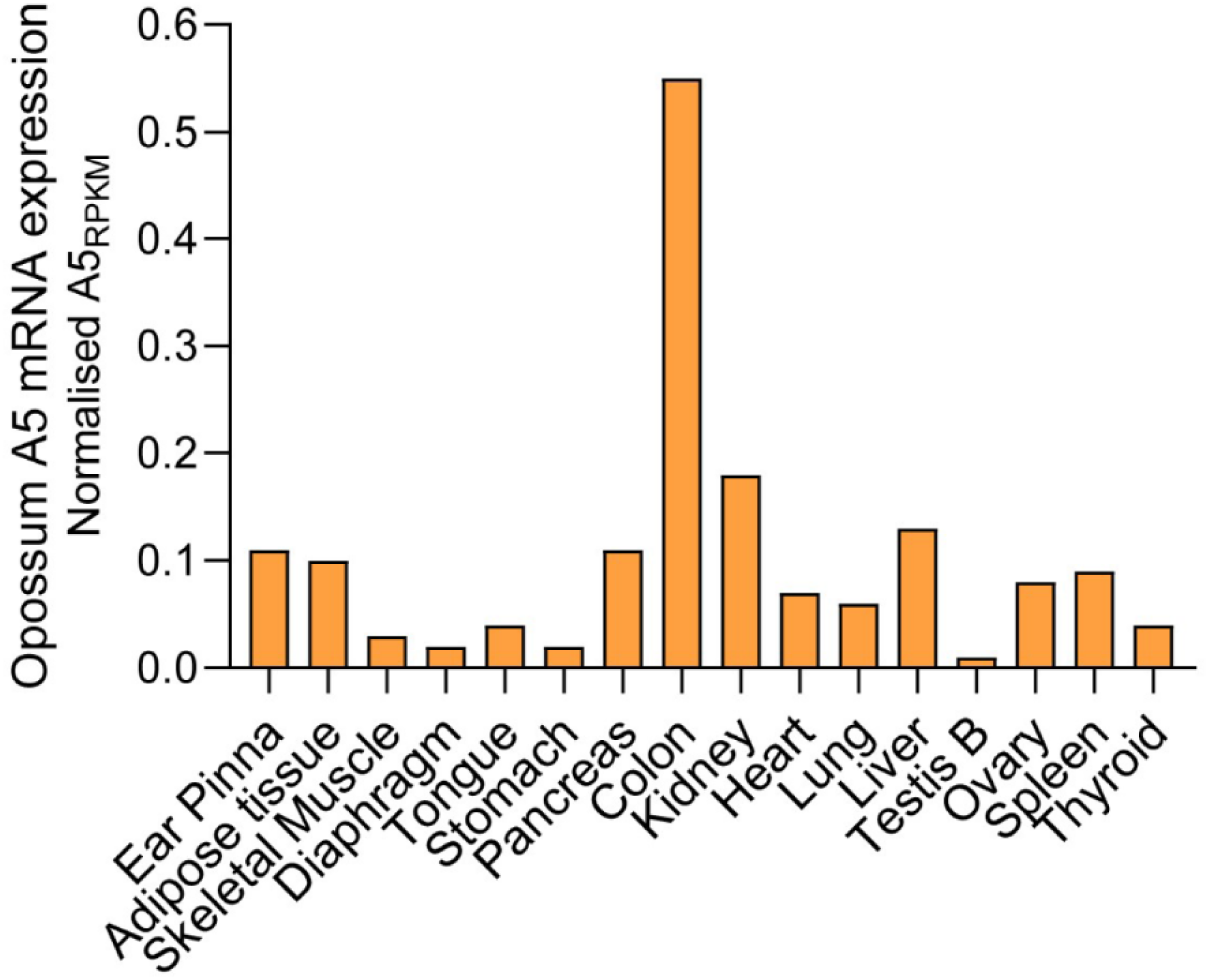
Relative expression of marsupial APOBEC5 (A5) across opossum tissues. Expression of A5, normalised to the housekeeping gene, RPS13, varies by tissue in the opossum.

APOBEC subcellular localisation and relative expression across different tissues may relate to its function. To begin assessing whether A5 is a marsupial functional homologue of placental A3, we analysed the intracellular localisation in mammalian cells of haemagglutinin (HA)-tagged A5 proteins from the Tasmanian devil (an Australian marsupial) and the gray short-tailed opossum (a South American marsupial). The rationale for studying A5 from these two species is that these Australian and American marsupials diverged at the root of extant marsupial lineages approximately 78 million years ago (mya) (Kumar, et al. 2022). Together, they represent the breadth of marsupial diversity, with Tasmanian devil A5 (tdA5) and opossum A5 (oA5) proteins sharing 72.4% amino acid sequence identity and highly similar predicted structures (Figure 2 and Supplementary Figure 1).

We compared the sub-cellular localisation of tdA5 and oA5 to human A3G (hA3G) in HEK293T cells by visual analysis of morphological differences in microscopic data. hA3G was included as a representative of the A3 family with strong antiviral activity (Figure 4A-C), as well as a mock transfection control (Supplementary Figure 4). hA3G primarily localised to the cytoplasm and concentrated in discrete mRNA processing bodies (P-bodies) as previously reported (Wichroski, et al. 2006; Stenglein, et al. 2008) (red arrows in Figure 4B). oA5 demonstrated broad subcellular distribution, localising to both the cytoplasm and the nucleus (Figure 4D-E, z-stack progression in Supplementary Video 1, and 3D projection in Supplementary Video 2), with apparent exclusion from nucleoli (pink arrows in Figure 4E). tdA5 showed similar cytoplasmic and nuclear localisation (Figure 4M-N, z-stack progression in Supplementary Video 3). oA5 rarely formed cytoplasmic puncta resembling P-bodies (yellow arrows in Figure 4H) but no examples of this were observed for tdA5. oA5 and tdA5 were also observed localising at the nuclear membrane (Figures 4J-K & 4P-Q, respectively, highlighted by white arrow in Figure 4J). Localisation in the cytoplasm may indicate an antiretroviral role for A5, while localisation in the nucleus suggests alternative/additional functions such as control of retrotransposons.

**Figure 4.**
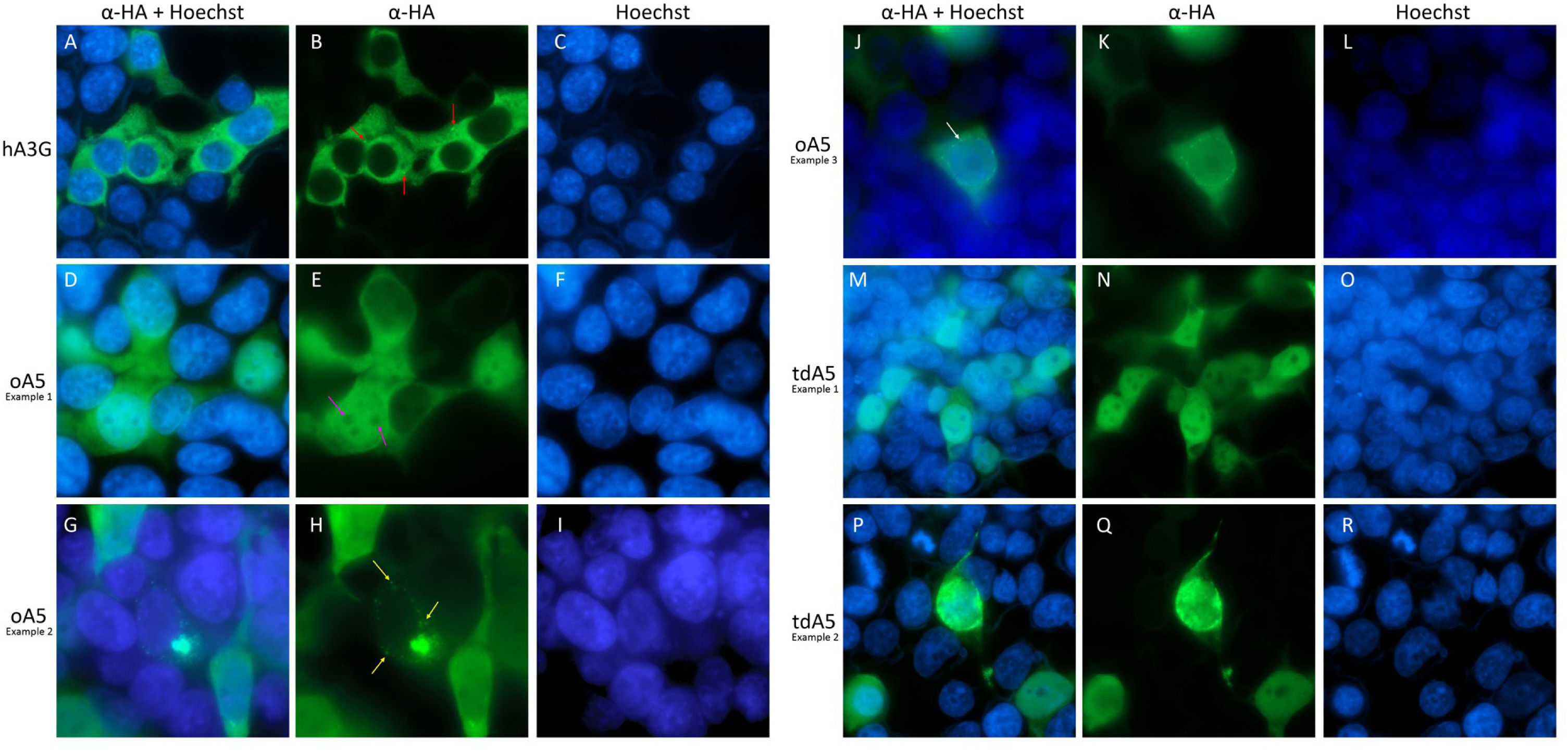
Subcellular localisation of APOBEC3 (A3) & APOBEC5 (A5) proteins in mammalian HEK293T cells. Haemagglutinin (HA)-tagged A3 and A5 proteins fluoresce green and Hoechst-stained nuclei fluoresce blue. All images were captured using a 63x1.4NA oil immersion lens. **A – C** Representative images of human APOBEC3G (hA3G) which localises primarily to the cytoplasm. Concentrations of hA3G in mRNA processing bodies (P-bodies) appear as bright green dots in the cytoplasm, indicated by the red arrows in B. **D – F** Representative images of opossum APOBEC5 (oA5) show a mixed distribution of oA5 in both the cytoplasm and the nucleus. Shadows representing nucleoli within the nucleus are indicated by the pink arrows in E. **G – H** Representative images of oA5 localising in concentrated spots within the cytoplasm that resemble P-bodies (yellow arrows). **J – L** Representative images of oA5 localising to the nuclear membrane indicated by the white arrow in J. **M – O** Representative images of Tasmanian devil APOBEC5 (tdA5) show a mixed distribution of tdA5 in both the cytoplasm and the nucleus. **P – R** Representation of a concentration of tdA5 at the nuclear membrane. Cells were stained with primary rabbit anti-HA C29F4 monoclonal antibodies (mAbs), secondary anti-rabbit Alexa Fluor 488 mAbs, and Hoechst 33342 nuclear stain.

### Marsupial A5 co-sediments with viral particles and is variably glycosylated

The antiviral role of placental mammalian A3 is dependent on packaging of A3 into viral particles that are released from the cell. When hA3G was co-expressed with an infectious molecular clone of murine leukemia virus (MLV) or Vif-deficient human immunodeficiency virus type 1 [HIVΔVif; Vif is the HIV-1 countermeasure to hA3G (Harris and Anderson 2016)] in HEK293T cells, it co-sedimented with viral particles in ultracentrifuged clarified cell culture supernatants as expected (Figure 5A). Similarly, oA5 and tdA5 co-sedimented with MLV and HIVΔVif viral particles (Figure 5A), which suggests it is incorporated into virions and has an antiviral function. oA5 and tdA5 are both 273 amino acids (AA) in length, sharing 72.4% amino acid sequence identity, and are predicted to have molecular masses of 31.7 and 32.1 kDa, respectively. Despite this size similarity, they were observed at markedly different positions relative to each other by SDS-PAGE under reducing conditions and Western blot analysis (Figure 5A). In contrast, when expressed in *Escherichia coli*, these proteins migrate, as expected, at the same position (Figure 6A). These data suggest that oA5 and tdA5 are variably glycosylated in mammalian cells, with oA5 being less glycosylated (or unglycosylated) compared to tdA5 (Figure 5A). oA5 and tdA5 sequences were analysed for predicted glycosylation sites using GlycoEP (Chauhan, et al. 2013). N-linked glycosylation sites were predicted for tdA5 amino acid positions 74 and 224, and for oA5 at positions 17, 79, 173, and 224. No O-linked glycosylation sites were predicted for either sequence. These predictions support the possibility that the different relative positions of oA5 and tdA5 observed by SDS-PAGE are due to glycosylation.

**Figure 5.**
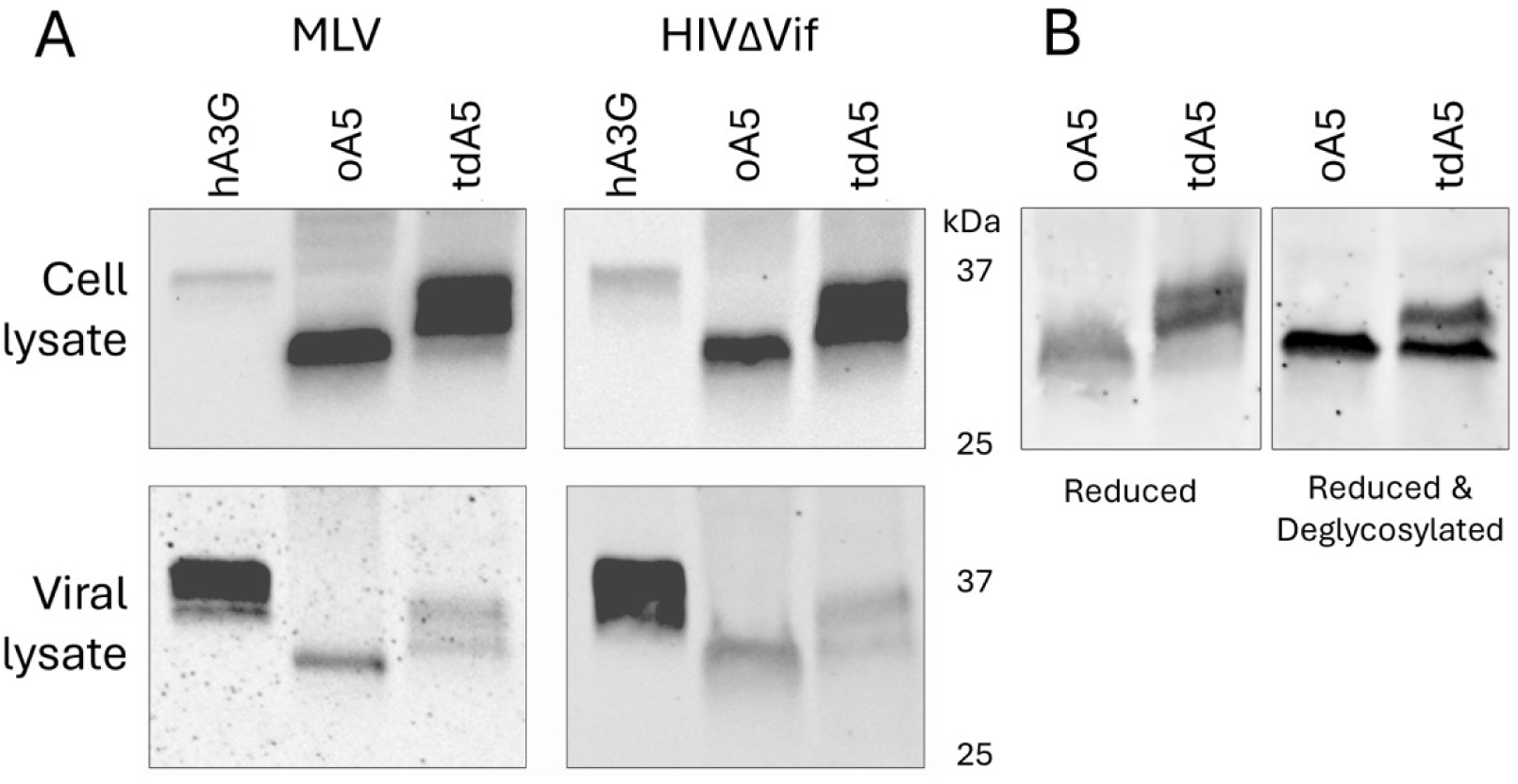
Marsupial APOBEC5 (A5) co-sedimentation in viral pellets, and glycosylation. **A** Human APOBEC3G (hA3G), opossum APOBEC5 (oA5), and Tasmanian devil APOBEC5 (tdA5) were expressed in HEK293T cells alongside the murine leukemia virus (MLV) or Vif-deficient human immunodeficiency virus (HIVΔVif). Viral particles were collected from clarified cell culture supernatant through ultracentrifugation through a 25% w/v sucrose cushion. Cells and viral pellets were lysed, samples were reduced and subjected to SDS-PAGE followed by Western blot analysis. **B** oA5 and tdA5 were expressed in HEK293T cells, and cell lysates were +/- treated with PNGase F to deglycosylate proteins, reduced and subjected to SDS-PAGE followed by Western blot analysis. Western blot was performed using primary rabbit anti-HA C29F4 monoclonal antibodies (mAbs), and secondary anti-rabbit Alexa Fluor 488 mAbs. Data are representative of N=3 independent assays.

**Figure 6.**
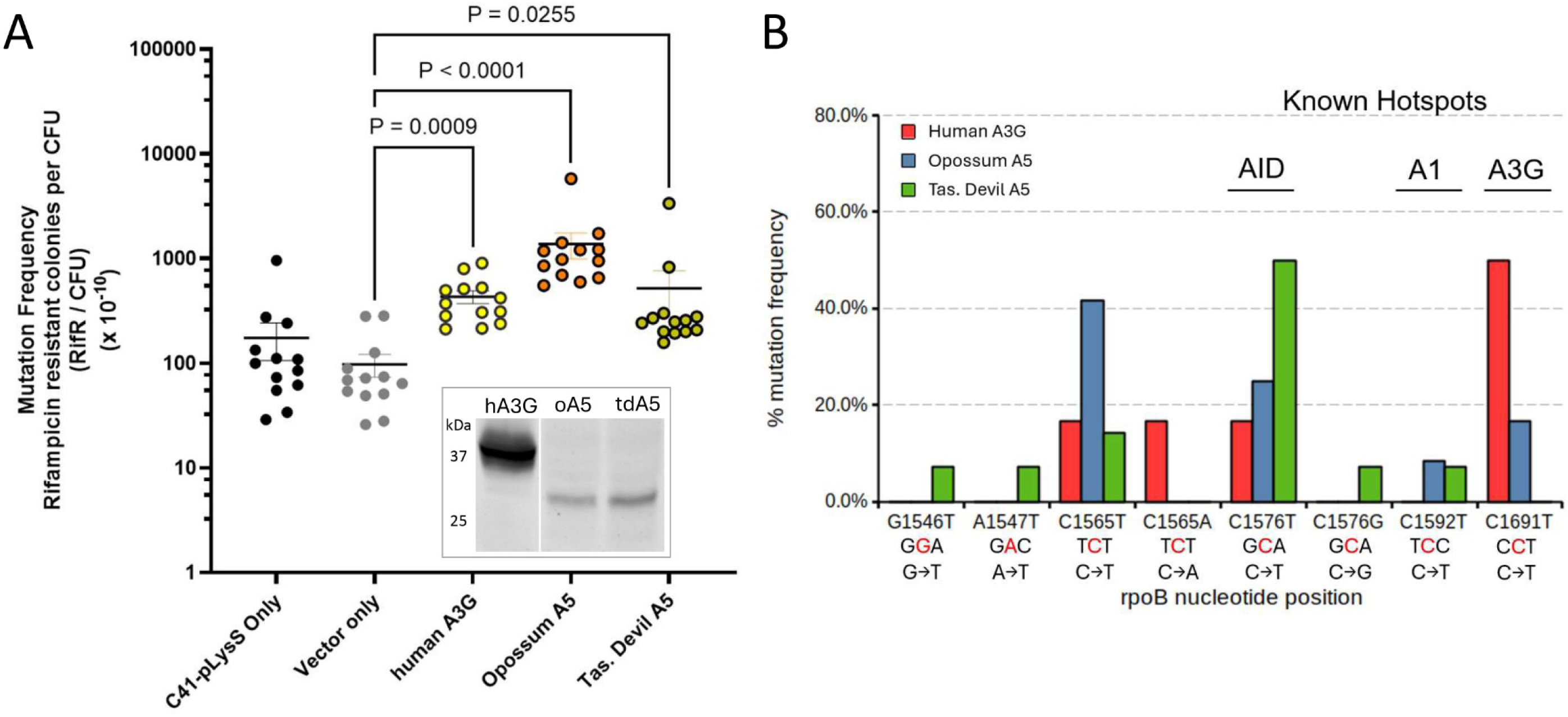
Functional analysis of DNA mutation by APOBEC5 (A5) proteins. **A** Haemagglutinin (HA)-tagged APOBEC3 (A3) and A5-expressing plasmids were transformed into C41-pLysS *E. coli* to measure mutation frequency through the emergence of rifampicin resistance. Each data point represents an independent sample and the median mutation frequency is represented by the bars. Test conditions were compared against untransformed *E. coli* (C41-pLysS only) and *E. coli* transformed with pET28a vector plasmid not encoding APOBEC (Vector only). Statistical significance was calculated by the Kruskal-Wallis test. **A-inset** Representative expression in *E. coli* of human APOBEC3G (hA3G), opossum APOBEC5 (oA5), and Tasmanian devil APOBEC5 (tdA5) proteins confirmed by Western blot/SDS-PAGE analysis. **B** Mutations elicited by hA3G, oA5, and tdA5 in a portion of the *rpoB* gene (nucleotide position 1,367 to 1,993 of the coding region) are revealed by Sanger sequencing of PCR amplicons. The identity and position of the mutated nucleotides are highlighted in red and indicated on the x-axis. Known mutation hotspots targeted by related AID/APOBEC proteins are indicated above. AID, Activation-induced deaminase; A1, APOBEC1.

To assess A5 glycosylation, we then subjected A5 proteins expressed in HEK293T cells to reducing and N-linked deglycosylating conditions in a subsequent SDS-PAGE/Western blot gel analysis. We observed that the position of the oA5 protein band was unchanged, indicating that oA5 was not deglycosylated (Figure 5B). tdA5 retained doublet bands, where the fastest migrating band shifted to a position matching PNGase-treated oA5 (Figure 5B). This indicates that tdA5 was glycosylated and that PNGase treatment either only partially deglycosylated tdA5 or that tdA5 contains additional post-translational modifications. Glycosylation of A5 may represent a unique protein modification among APOBEC family members.

### Marsupial A5 is a DNA mutator

To evaluate if A5 is capable of mutating DNA, we employed a rifampicin mutagenesis assay that has been previously used to evaluate AID/APOBEC family protein function (Harris, et al. 2002; Hayward, et al. 2018), using hA3G as a positive control. Mutations in the *E. coli rpoB* gene, which generate antibiotic resistance to rifampicin, increase with APOBEC-induced mutagenic stress (Harris, et al. 2002; Shi, et al. 2015). We observed mutation frequencies above the background level for both oA5 and tdA5, with oA5 eliciting a 10.3-fold (p<0.0001, N=12) greater mutation frequency than vector only (Figure 6A).

Different AID/APOBEC proteins are known to display bias in the specific mutations they induce in *E. coli rpoB*, represented as mutation ‘hotspots’. Sanger sequencing of the *rpoB* gene from individual colonies revealed the presence of specific mutations (Figure 6B). The hotspots for oA5 and tdA5 were at C1565 and C1576, respectively (Figure 6B). These sites have previously been reported as APOBEC targets, and C1576 is a hotspot for AID (Harris, et al. 2002). These data demonstrate that A5 is catalytically active and capable of mutating DNA, and that different A5 proteins have distinct preferences in the sites that they target.

### Marsupial A5 restricts HIV-1, but not MLV, and is counteracted by HIV-1 Vif

APOBEC proteins exert their inhibitory effect by being packaged into viral particles in the producer cell and interfering with viral replication in the target cell (Cullen 2006). To determine whether marsupial A5 has a functional antiviral role homologous to placental A3, we first assessed its impact on the infectivity of two model retroviruses, MLV (genus *Gammaretrovirus*) and a version of HIV-1 lacking the A3 countermeasure, Vif (HIVΔVif; genus *Lentivirus*), both of which are restricted by placental A3 proteins (Kobayashi, et al. 2004; Rulli, et al. 2008; Albin and Harris 2010). hA3G was included as a positive control that is known to restrict HIVΔVif and MLV (Rulli, et al. 2008).

To test the antiviral activity of A5, cells were co-transfected with plasmids encoding MLV or HIVΔVif and increasing concentrations of plasmids encoding oA5 or tdA5. The antiviral activity was compared to the infectivity of virus generated without the addition of any APOBEC. Cell culture supernatants containing viral particles were collected and clarified, then normalised for viral particle content. Normalised supernatants were then used to infect mouse NIH/3T3 cells with MLV, or human TZM-bl cells with HIVΔVif. Infection with MLV and HIVΔVif was measured by virion-associated reverse transcriptase (RT) activity in NIH/3T3 cell culture supernatants and by luciferase activity in TZM-bl cell lysates, respectively.

Infectivity of both MLV and HIVΔVif was strongly inhibited by hA3G, with a reduction of 90% inhibition of HIVΔVif (Figure 7A) and 97% inhibition of MLV (Figure 7C). oA5 and tdA5 modestly inhibited HIVΔVif (one-way ANOVA P=0.007 and P=0.001, respectively), with both reducing infectivity by approximately 40% at the same dose as hA3G, with an infectivity trend reflecting a dose-response (linear trend slopes of -13.42 for oA5 [R^2^=0.99, P=0.0009] and -11.67 for tdA5 [R^2^=0.79, P=0.0004]). While both oA5 and tdA5 appeared to have a minor impact on the infectivity of MLV (one-way ANOVA P-values of 0.033 and 0.073, respectively), a linear dose-response was not observed for oA5 and was weak for tdA5 (linear trend slopes of -0.01 for oA5 [R2=0.01, P=0.71] and -0.05 for tdA5 [R2=0.74, P=0.024]). These data suggest that HIVΔVif is modestly inhibited by A5 while MLV is largely resistant to inhibition by A5.

**Figure 7.**
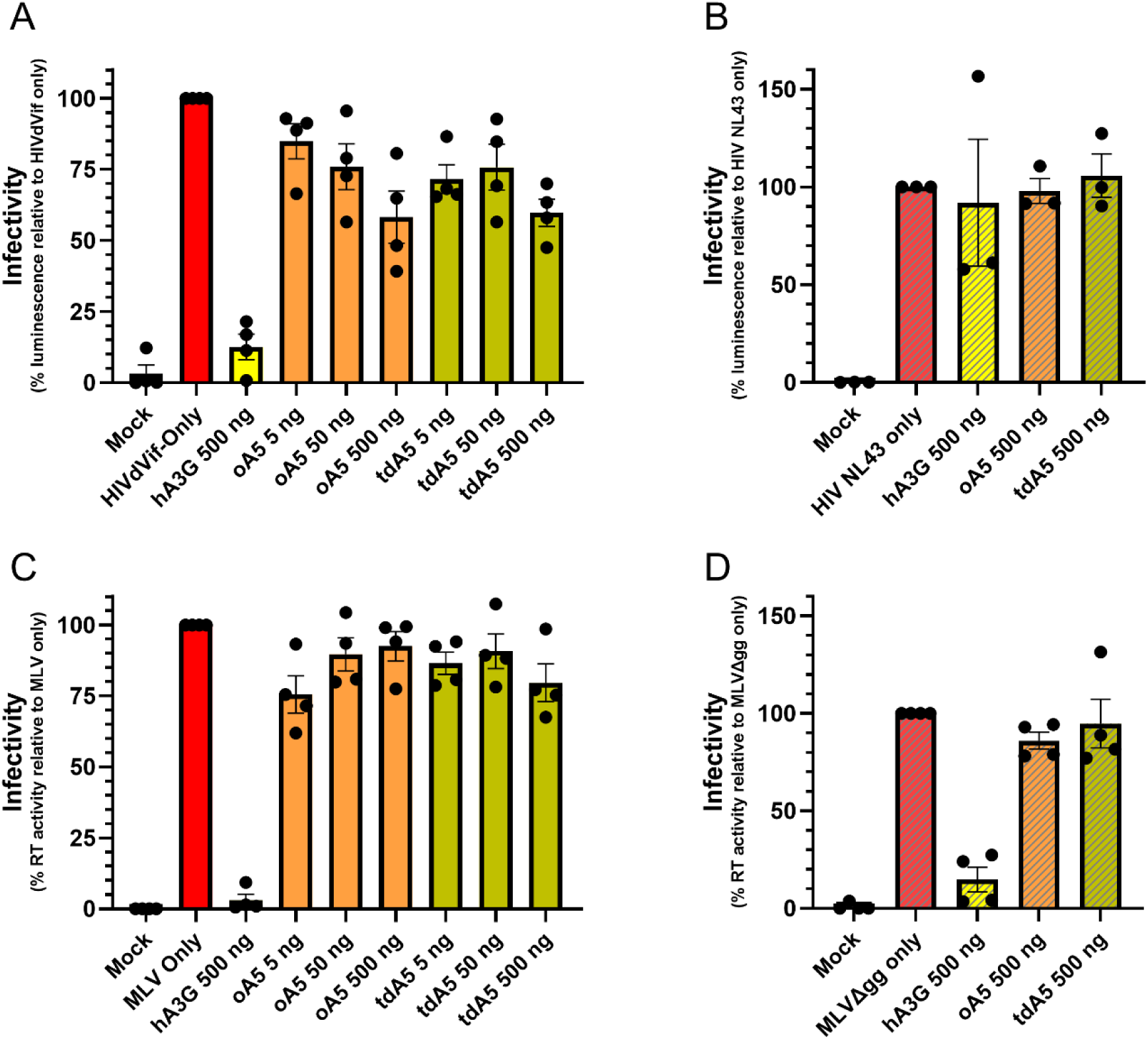
APOBEC5 (A5)-mediated inhibition of infectivity of HIV, but not MLV. **A** Vif-deficient human immunodeficiency virus type 1 (HIVΔVif), **B** Vif-competent HIV (HIV NL4.3), **C** murine leukemia virus (MLV), or **D** glyco-Gag-deficient MLV (MLVΔgg) viral particles were generated in HEK293T cells in the presence of human APOBEC3G (hA3G), opossum APOBEC5 (oA5), and Tasmanian devil (tdA5) proteins to assess the inhibitory effect of A5 on viral particle infectivity. Cells were co-transfected with plasmids expressing HIVΔVif, HIV NL4.3, MLV, or MLVΔgg and variable concentrations (5-500 ng) of A3 or A5 expression plasmids. HEK293T cell culture supernatants were normalised for viral particles by virion-associated reverse transcriptase activity. Human TZM-bl cells or mouse NIH/3T3 cells were inoculated with normalised HIVΔVif / HIV NL4.3 and MLV / MLVΔgg, respectively. Infectivity relative to HIVΔVif, HIV NL4.3, MLV or MLVΔgg alone was measured by quantification of luciferase activity in TZM-bl cell lysates or virion-associated reverse transcriptase activity in NIH/3T3 cell culture supernatants. Statistical significance was evaluated by one-way ANOVA (**A** & **C**) and paired t-tests (**B** & **D**). Error bars represent the SEM of N=3-4 independent assays.

We next addressed if the modest restriction of HIV could be counteracted by Vif, and if the lack of restriction of MLV was due to the glyco-Gag isoform of MLV gag, which counteracts restriction by murine A3 (Rulli, et al. 2008; Stavrou, et al. 2013). To test these questions, we performed the A3/A5 co-transfections using a Vif-competent version of HIV (HIV NL4.3) and a glyco-Gag-deficient version of MLV (MLVΔgg). Both HIV NL4.3 and MLVΔgg were not restricted by A5 (Figures 7B and 7D). These data indicate that the presence of Vif counteracts restriction by A5, and that the lack of restriction of MLV is not a due to counteraction by glyco-Gag.

### Marsupial ERVs reveal differential APOBEC-like restriction of recently endogenised retroviruses

To determine if modern marsupials contain inherited genomic signatures of A3-like mutation of ancient retroviral infections, we performed a hypermutation analysis of groups of endogenous gammaretroviruses and betaretroviruses in the genome of the gray short-tailed opossum. Many of the analysed ERVs were intact, with one of eight gammaretroviral ERVs and six of ten betaretroviral ERVs containing no frameshift or premature stop codon mutations. For all ERVs, analysis of the target site duplications confirmed that each ERV represented a unique infection/integration event (Supplementary Table 2). Molecular clock analyses determined that the gammaretrovirus and betaretrovirus groups had infected the opossum germline between <0.40 – 4.27 and <0.46 – 2.31 mya, respectively (Supplementary Table 2). The sequences of each opossum ERV used in this analysis are provided as Supplementary Data 1.

A3 activity mutates C-to-U in the negative strand of DNA during retroviral reverse transcription, leading to a bias toward positive strand G-to-A mutations in ERVs in the host genome (Perez-Caballero, et al. 2008). A3 activity also depends on local sequence context. The presence of context-specific, strand-biased mutagenic signatures in ERVs constitutes definitive evidence of interaction between retroviruses and A3 and/or A3-like enzymes (Lee, et al. 2008).

Phylogenies for each group of ERVs were estimated with the maximum likelihood method and ancestral sequence reconstruction was undertaken by calculation of the common ancestor of each group (Supplementary Figure 5). The mutations in each ERV were compared against each group’s common ancestral sequence. Strand-biased G-to-A hypermutation with dinucleotide context bias, predominantly GY (GT or GC), was identified in gammaretroviruses (Figure 8A-B). No evidence of hypermutation was identified in betaretroviruses (Figure 8C-D). These data indicate that although marsupials do not possess *A3* genes, other members of the marsupial APOBEC repertoire function as antiviral restriction factors in an A3-like manner, against some but not all retroviruses.

**Figure 8.**
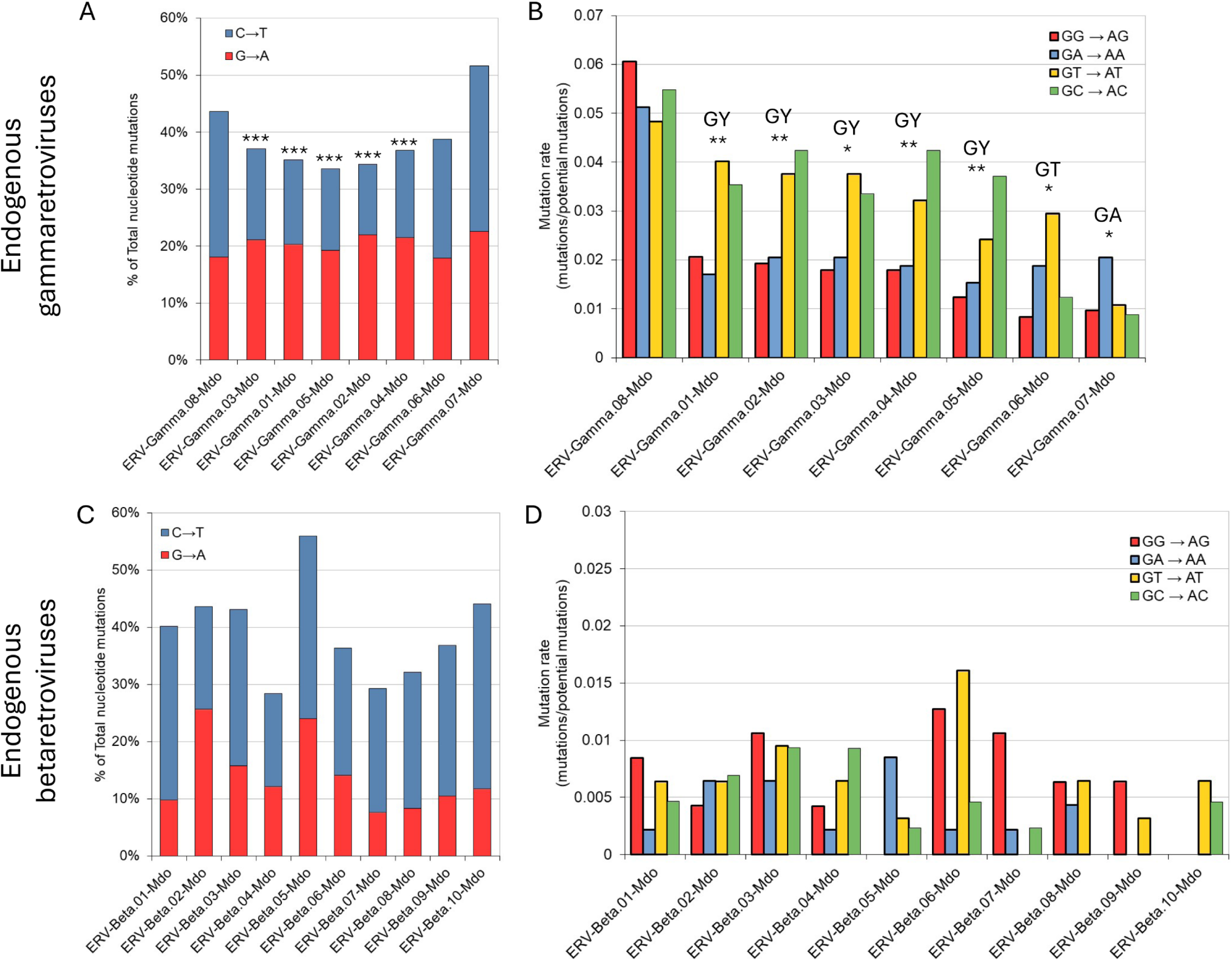
Hypermutation analysis of endogenous retroviruses (ERVs) in the opossum genome reveals signatures of A3-like activity in gammaretroviruses but not betaretroviruses A &. **C** Comparison of G-to-A versus C-to-T mutations on the positive coding strand of ERVs reveals strand-biased cytosine deamination in gammaretroviruses (A) but not betaretroviruses (C). **B & D** Analysis of the context of coding strand G-to-A mutatons reveals a GY (GT or GC) dinucleotide bias in gammaretroviruses (B) but not betaretroviruses (D). Fisher’s Exact Test was used to calculate statistical significance (*P<0.05; **P<0.01; ***P<0.001).

## Discussion

In this study, we investigated A5, a unique and previously uncharacterised member of the APOBEC family, to determine its evolutionary history, structure, and potential function. Using BLAST homology searches, we identified A5 homologues in marsupials, reptiles, amphibians, and jawed fish, but not in birds, monotremes, or placental mammals, suggesting lineage-specific gene loss. Phylogenetic and structural analyses confirmed A5’s classification within the APOBEC family, with conserved cytosine deaminase motifs and a predicted structure similar to other members of the AID/APOBEC family. Within marsupial tissues we found broad expression, which was highest in the colon, suggesting a possible multifunctional role. Cellular localization experiments revealed that ectopically expressed marsupial A5 is present in both the cytoplasm and nucleus. Functional assays demonstrated that A5 can mutate DNA and restrict the infectivity of the retrovirus HIVΔVif, which lacks the APOBEC3 countermeasure Vif, and that A5 is counteracted by Vif. Additionally, glycosylation analysis suggested that A5 undergoes post-translational modifications, a feature not previously reported in APOBEC proteins to our knowledge. Finally, evolutionary analysis of marsupial ERVs revealed signatures of A3-like antiviral activity, indicating that A5 or another marsupial APOBEC protein restricts retroviral infections. These findings support the hypothesis that A5 is a conserved, functional cytosine deaminase and that APOBEC proteins have played a role in marsupial antiviral defence.

### A5 sequences are common to jawed vertebrates and have been lost from several lineages

A5 is a novel, putative member of the AID/APOBEC protein family whose existence has previously only been cursorily reported based on the homology and phylogenetic relatedness of a limited number of APOBEC-like sequences (1x amphibian, 1x reptile, 4x marsupial) to known APOBEC proteins (Severi, et al. 2011; Ito, et al. 2020). An important aspect of *A5* is its absence from the genomes of placental mammals. The original report of the existence of A5, in 2011 (Severi, et al. 2011), was made when the number of sequenced animal genomes was far fewer than are available today.

To improve our understanding of the taxa which do or do not possess this novel APOBEC, we searched for sequences homologous to previously reported A5 sequences across all vertebrate taxa represented in GenBank. A5 homologues were identified in 36 species of marsupial mammals, reptiles, amphibians, and jawed fish (Figure 1). The presence of A5 in jawed fish is a novel finding and indicates that it was present in the last common ancestor of all jawed vertebrates, as is the case for A2 and A4, making A5 a more ancient APOBEC than A3, which is only present in placental mammals, and A1, which is only present in amniote vertebrates (Salter, et al. 2016). It also reveals that A5 has been lost in several key vertebrate lineages including birds, monotremes, and placental mammals. Because marsupials diverged from placental mammals after the divergence from monotremes, this means that *A5* genes were lost from placental and monotreme mammals in independent events. Together with its absence in birds, this reveals that *A5* has been lost on at least three separate occasions.

It might be reasonable to speculate that the loss of *A5* in placental mammals may have been due to some redundancy of function provided by the emergence of *A3*, but this does not appear to be the case for monotremes or birds, which lack *A3* and have not been reported to possess other novel APOBECs. Three of the fish species in which A5 was found are different sturgeon species in which a duplication of *A5* has occurred (Figure 1B). This appears to be the result of a duplication in their common ancestor as the A5A of each species was more similar to the A5A of the other species (95-96% amino acid identity) than it was to A5B within the same species (91-93% amino acid identity), and each group clustered phylogenetically (Figure 1B). Such duplication events are common among mammalian *A3* genes (Münk, et al. 2012; Hayward, et al. 2018) and may reflect diversification of function or an increase in the range of proteins that they can interact with (or avoid interaction with), such as might be the case when interacting with viral targets and countermeasures.

### A5 possesses conserved APOBEC motifs and structure, and are uniquely glycosylated

A5 has retained the HxEx_y_PCx_2-5_C Z-domain required for cytosine deaminase activity and the predicted N-terminal structure of A5 proteins is highly similar to placental A3 proteins (Figure 1C and 2A-C), possessing the canonical cytosine deaminase fold (Salter, et al. 2016), despite the difference in primary amino acid sequences. For example, Tasmanian devil A5 and human A3A share only 24.5% amino acid sequence identity across their aligned N-terminal region which contains the Z-domain. This indicates that for APOBEC proteins, the conservation of structure, along with a handful of specific residues critical for catalytic activity, is more important than conservation of the bulk of the primary amino acid sequence. This has been observed for other innate immune proteins, such as tetherin (BST-2), which are similarly subject to an ‘arms race’ with viral antagonists (Perez-Caballero, et al. 2009; Blanco-Melo, et al. 2016; Hayward, et al. 2022). A5 proteins were found to have a unique, extended C-terminal region that does not share homology to other APOBEC proteins and is not predicted to contain an intrinsically stable structure (Figure 2A-B, Supplementary Figure 1). One other member of the AID/APOBEC family, A4, similarly contains an extended C-terminal region whose function has not been reported but is also predicted to be intrinsically disordered (Salter, et al. 2016) (Supplementary Figures 2 & 3). This could be taken to indicate some similarity of mechanism between A4 and A5.

Disordered regions in proteins can be involved in functions such as signalling, localisation, or protein-protein interactions (Wright and Dyson 1999), which in the case of APOBECs, might involve substrate specificity or interactions with viral targets and countermeasures. For example, A2 has an N-terminal disordered region hypothesised to influence its subcellular localisation (Pecori, et al. 2022). The presence of protein motifs, 239PxK and 256KRKL in A5 (Figure 2E) at the extremity of the C-terminal region, which are highly conserved across all taxa, suggest a critical function for this domain.

Surprisingly, we found that Tasmanian devil A5 was glycosylated (Figure 5B). Glycosylation is a common post-translational modification critical to the biological functions of many proteins, but to our knowledge has not been reported for any AID/APOBEC protein. It is curious that we did not observe glycosylation for opossum A5, and this difference suggests a potential diversification in the function or activity of A5 across marsupial taxa. Australian and American marsupials, represented in this study by the Tasmanian devil and gray short-tailed opossum, are distantly related, with their last common ancestor placed at the root of all extant marsupial lineages approximately 78 mya (Kumar, et al. 2022). Given the distance of their relationship, it is perhaps unsurprising that their A5 proteins have differentiated, potentially reflecting evolutionary pressures unique to each lineage. In the context of APOBEC proteins, it is possible that glycosylation could impact protein-protein interactions. A prominent example of this is glycosylation of the Gag protein of the retrovirus, MLV (Rosales Gerpe, et al. 2015). Variation in glycosylation of MLV Gag modulates the interaction of murine A3 (mA3) with MLV Gag, balancing the restrictive activity of mA3 with a beneficial boost to MLV diversification through mA3-induced mutagenesis (Rosales Gerpe, et al. 2015). In the context of a hypothetical antiviral role for A5, glycosylation may enable binding to viral targets or prevent binding by viral countermeasures.

### A5 has broad subcellular localisation and tissue expression suggesting a possible multifunctional role

We assessed the subcellular distribution of marsupial A5 to gain insight into possible A5 functions. A5 was seen to be broadly distributed in both the cytoplasm and the nucleus (Figure 4). Within the nucleus, A5 appeared to be largely excluded from nucleoli (e.g., Figure 4E). A5 was occasionally observed concentrated at the nuclear membrane (e.g., Figure 4J-K) and rarely in cytoplasmic spots resembling ‘P-bodies’ (Wichroski, et al. 2006) (e.g., Figure 4H). In some cells, A5 was seen to localise mostly in the cytoplasm while in others it was seen mostly in the nucleus (examples of both in Figure 4E) which may indicate that localisation varies during different phases of the cell cycle, as has been reported for various A3 proteins (Lackey, et al. 2013). APOBEC proteins that have a primary role in activities such as antiviral immunity (e.g., hA3G), or RNA editing (e.g., A1), localise mainly to the cytoplasm (Salter, et al. 2016). APOBECs that localise to the nucleus are involved in mutating nuclear DNA with roles such as antibody development (e.g., AID) or cancer mutagenesis (e.g., A3B) (Salter, et al. 2016). A presence in both nuclear and cytoplasmic locations suggests that A5 may have multiple roles. Other possibilities exist, for example A3C is distributed to both the cytoplasm and nucleus and plays a role in inhibition of retrotransposon activity, and its anti-retrotransposon activity is not dependent on localisation (Muckenfuss, et al. 2006).

A5 proteins were also found to co-sediment with viral particles collected from cell culture supernatants subjected to ultracentrifugation (Figure 5A). This suggests that A5 is incorporated into virions in the same manner as antiviral A3 proteins (Cullen 2006), with the caveat that A5 may be present by other mechanisms, such as being present in non-viral extracellular vesicles that have co-sedimented with the virus.

Biased tissue expression of APOBEC proteins can also be indicative of their function. For example, mammalian A1 is predominantly expressed in the small intestine, correlating with its role in lipid metabolism (Nakamuta, et al. 1995). AID is highly expressed in immune tissues and specifically B-cells for its role in antibody diversification (Muto, et al. 2000), and various A3 are expressed in immune cells and tissues where they can encounter their viral targets (Koning, et al. 2009). Analysis of A5 mRNA expression across a multi-tissue transcriptome of the gray short-tailed opossum provides clues to the role of A5 (Figure 3; GenBank: PRJNA200320). A5 was broadly and variably expressed across all tissues, but with a starkly higher expression in the colon than any other analysed tissue (Figure 3), followed by the liver and kidneys. Expression of APOBEC proteins in the colon and liver is linked with roles including RNA editing, lipid metabolism, and cancer mutagenesis (Huang, et al. 2010; Blanc, et al. 2014; Lan, et al. 2014), and expression in the kidneys is linked with viral pathogenesis (Peretti, et al. 2018) and pro-survival responses to kidney damage (Guo, et al. 2023).

That the highest expression was in the colon could suggest a possible role involving the gut microbiome, and we hypothesise that a colon-enriched APOBEC could function in antiviral defence against gut viruses or modulating host RNA stability (in a manner similar to APOBEC1) to regulate gene expression in response to microbial metabolites. That we also observe modest expression, relative to the colon, across most other tissues might indicate a broader, more generalised function such as the control of retrotransposon activity. These are not mutually exclusive possibilities. A3 proteins for example, act in both antiviral defence and genome stability across tissues through control of retrotransposon activity (Koito and Ikeda 2013; Friedli and Trono 2015; Modenini, et al. 2022). These findings suggest that a similarly multifunctional role may be played by marsupial A5.

Regarding a possible antiviral role for A5, retroviruses infecting marsupials are known to be detectable in faeces and urine, which respectively pass through the colon and kidneys, where the second highest expression was observed (Figure 3) (Kawakami, et al. 1977; Hayward, et al. 2020). It is also worth noting that there are multiple important structural and physiological differences between the colons of marsupials and placental mammals pertinent to gene expression, with herbivorous marsupials (e.g., brush-tailed possum) possessing a larger, more complex colon that is involved in hind-gut fermentation and the metabolism of volatile fatty acids (Hume 1989; Leng 2018). However, the gray short-tailed opossum analysed here is an omnivorous, leaning toward carnivorous, marsupial with a short colon more similar to small placental carnivores/insectivores (Santori, et al. 2004; de Carvalho, et al. 2019). As such, the function of A5 is unlikely to be confined to a role related to the differences in marsupial vs. placental colon physiology.

### Marsupial A5 mutates DNA and modestly restricts the infectivity of a model retrovirus

Assessment of the function of A5 using a rifampicin mutagenesis assay demonstrated that both opossum and Tasmanian devil A5 were capable of mutating DNA when expressed in *E. coli*, causing an increase in the emergence of rifampicin resistance through the generation of mutations in the *E. coli rpoB* gene (Figure 6A). oA5 had a greater mutagenic effect than hA3G, and both oA5 and tdA5 elicited mutations at discrete hotspots within *rpoB* (Figure 6B). This is consistent with the DNA editing functions of other AID/APOBEC proteins and confirms that A5 can function as a mutagenic cytosine deaminase. This is an important differentiating factor, because some members of the AID/APOBEC family, specifically APOBEC2 and APOBEC4, appear to be incapable of catalytic deamination activity (Harris, et al. 2002; Mikl, et al. 2005; Lada, et al. 2011).

To evaluate the potential of A5 to act as an antiviral restriction factor we assessed its ability to restrict the infectivity of two model retroviruses, MLV and a Vif-deficient clone of HIV-1, HIVΔVif. When HIVΔVif was expressed in the presence of oA5 and tdA5 in the producer cell, we observed a modest, dose-dependent reduction in viral particle infectivity (Figure 7A). This effect was not observed when oA5 and tdA5 were expressed in the presence of a Vif-competent clone of HIV-1, HIV NL4.3 (Figure 7B), which might indicate that A5 is counteracted in a similar manner to placental A3. MLV particles expressed in the presence of A5 were not observed to have a robust dose-dependent impact on their infectivity (Figure 7C). Glyco-Gag is a glycosylated isoform of the MLV Gag protein and unique inhibitor of mouse A3, which does not prevent packaging of mouse A3 into virions but instead prevents access to the reverse transcription complex in the viral core (Rulli, et al. 2008; Stavrou, et al. 2013). We suspected that glyco-Gag might act as a countermeasure to A5 in a similar manner to A3 because A5 appeared to be packaged into MLV virions (Figure 5A) but no restriction was observed (Figure 7C). However, when we evaluated the effect of A5 against a version of MLV that was not capable of expressing glyco-Gag, MLV remained unrestricted (Figure 7D), indicating that its resistance to restriction was not due to glyco-Gag. Together, these results indicate that A5 may function as an antiviral restriction factor, which is susceptible to counteraction by viral proteins such as HIV-1 Vif, and by an unknown mechanism in the case of MLV.

A3 proteins have been demonstrated to restrict retroviral replication by two important mechanisms, (i) deamination-dependent hypermutation of viral cDNA (Harris, et al. 2003), and (ii) deamination-independent stearic hindrance of the synthesis of viral cDNA (Belanger, et al. 2013). While A5 is a capable of DNA mutation, ongoing studies should determine whether or not the restrictive effect against HIV-1 seen here is deamination-dependent.

### Naturally occurring retroviral insertions in the opossum show evidence of differential A3-like restriction

While APOBEC-mediated restriction of retroviral infection may be observable in vitro, it is not a given that experimental over-expression systems reflect true biological functions. APOBEC1 for example, has been reported to restrict retroviral replication in vitro, but follow-up studies using A1 knock-out mice have revealed that this function is not observed in vivo (Petit, et al. 2009; Barrett, et al. 2014). An essential component of retroviral replication is the integration of the provirus into the genome of the infected cell, which occasionally occurs in gamete cells. When the latter happens within gametes that generate offspring, the retrovirus becomes a heritable genetic element, generating a genetic ‘fossil’ record of ancient infections in the genomes of host species (Johnson 2019). As a result, it is possible to analyse endogenous retroviruses in animal genomes for signatures of A3/A3-like mutagenic activity (Lee, et al. 2008; Hayward, et al. 2018).

In general, marsupial genomes have been invaded by retroviruses to a lesser degree than placental mammals (Ito, et al. 2020; Harding, et al. 2024). For example, ∼8% of the human genome is derived from retroviruses, while Australian marsupials have less than 1% retroviral content, with Dasyuridae (including Tasmanian devils) having the greatest amount at 0.72% (Ito, et al. 2020; Harding, et al. 2024). The gray short-tailed opossum stands in stark contrast, with a retroviral content of approximately 10%, and is second only to some rodents for total retroviral content (Ito, et al. 2020). It has previously been reported that diverse vertebrate genomes, including marsupials, contain evidence of APOBEC-like G-to-A positive strand biased DNA editing activity in retrotransposons (Knisbacher and Levanon 2016). To expand on this, we looked for A3-like mutational signatures in a representative marsupial genome (the gray short-tailed opossum), including critical dinucleotide contextual biases, relative to their unintegrated ancestral viral sequence, reflecting an antiviral immune response to infection with an exogenous retrovirus. We inferred the ancestral sequences of groups of recently integrated (<0.4 – 4.3 mya) gammaretroviruses and betaretroviruses (Figure 8). Both of these groups contain multiple fully intact ERVs, which means they may represent extant infectious retroviruses and are potentially capable of generating infectious viral particles. However, this would need to be experimentally confirmed as there are difficulties involved in differentiating between ERVs capable and incapable of generating exogenous viral particles by their sequence alone (Hayward and Tachedjian 2021).

Intriguingly, we found an A3-like signature with G-to-A positive strand-bias and a specific GY-to- AY dinucleotide context across the gammaretroviral ERVs but no evidence of A3-like restriction in the betaretroviral ERVs (Figure 8). The GY (GC or GT) dinucleotide preference is a point of distinction from A3 proteins, which have GR (GG or GA) biases (Salter, et al. 2016). The hypermutation of gammaretroviruses with a consistent dinucleotide context bias, relative to the ancestral retroviral sequence is strongly indicative of A3-like antiviral restriction. An important caveat is that this signature is not necessarily the result of activity by A5, as other marsupial APOBEC proteins, such as A1, may also be involved. Importantly, the fact that no A3-like signatures were seen in the betaretrovirus group may be an indication of the presence of a betaretroviral countermeasure against marsupial APOBEC proteins. This has been observed in betaretroviruses infecting primates (Doehle, et al. 2006), the existence of which would further validate a role for marsupial APOBECs as antiviral restriction factors.

Opossum A1 has been reported to have modest antiretroviral activity in vitro, to a similar extent as demonstrated for A5 in this study, and showed similar patterns of cell localisation and widespread tissue expression (Ikeda, et al. 2017). Whether opossum A1 or A5 act as antiviral restriction factors in vivo remains to be determined, and which of these APOBECs are responsible for the A3-like signature in opossum gammaretroviruses is unknown. It may be the case that both are responsible for antiviral restriction in natural infections. Certainly, the case of expansion of antiviral A3 in placental mammals makes it clear that there is often the need for multiple antiviral APOBEC proteins to be present. These findings suggest a case of convergent evolution where A5 and/or A1 play a similar role in marsupials that A3 plays in placental mammals and highlight the importance of APOBEC-mediated antiviral activity.

### Limitations and next steps

Here we presented an important first step in the characterisation of an understudied member of a protein family involved in essential biological functions and antiviral defence. Naturally, many questions remain to be answered. Regarding the antiviral function of A5, it will be a priority to establish whether or not the modest retroviral restriction observed is deamination-dependent with follow-up mutagenesis studies to shed light on the A5 mechanism of action. Although we tested oA5 and tdA5 against model retroviruses, we did not assess activity against species-specific exogenous retroviruses, which have yet to be identified in opossums or Tasmanian devils. Should any ERVs from these hosts be found capable of generating infectious viral particles, A5’s antiviral activity against them could be evaluated.

Our observation that Tasmanian devil A5 but not opossum A5 is glycosylated suggests a diversification of A5 function among marsupials. Ongoing work should assess if glycosylation influences stability, localization, or viral interactions. The conserved, intrinsically disordered C-terminal tail across taxa suggests a crucial, yet uncharacterized function, and future analyses should aim to determine its function, possibly in cellular targeting or binding.

While we focused on antiviral activity, A5 may have broader roles, such as retrotransposon control or RNA-editing, given its nuclear and cytoplasmic subcellular distribution. Its high expression in the opossum colon also raises the possibility of gut-related functions, which could be explored through transcriptomics and metabolomics. With gene knockout models available in *Monodelphis domestica*, functional studies of marsupial APOBECs are now possible (Kiyonari, et al. 2021).

## Conclusions

This study represents the first report on the characterisation of A5, a novel structurally unique member of the AID/APOBEC protein family that was present in the last common ancestor of all jawed vertebrates which has been lost from multiple extant taxa. A5 is catalytically active and capable of deaminating DNA, has broad subcellular and tissue distribution, and displays antiviral activity against a model retrovirus, HIV-1, which is counteracted by HIV Vif. Ancient retroviral fossils in the opossum genome contain unambiguous evidence of differential A3-like activity against gammaretroviruses, revealing that even though marsupials lack A3, a homologous antiviral function is fulfilled by other APOBEC proteins. These findings shed new light on convergent evolution in the innate immune system between marsupial and placental mammals.

## Methods

### Identification of A5 sequences

To identify sequences homologous to previously reported A5 proteins, we searched the publicly accessible GenBank records using the protein basic local alignment search tool (BLAST; blastp) portal (https://blast.ncbi.nlm.nih.gov/Blast.cgi). The search query was the Tasmanian devil A5 amino acid sequence (GenBank: XP_003765824), with expect value set to <1x10^-30^, max target sequences set to 1000, and other parameters set to default values.

### Phylogenetic and sequence analyses

To assess how the identified A5 sequences relate to other AID/APOBEC proteins, we created a multiple sequence alignment using MUSCLE (Edgar 2004). This alignment included all A5 sequences, along with selected sequences from each AID/APOBEC group to represent their presence across different species. We also included CDA as an outgroup. The phylogeny was generated using the maximum likelihood method in CLC Genomics v24.0 (CLC, QIAGEN, Venlo, Netherlands) with the best-fit JTT+G+I model and 1,000 bootstrap replicates. The tree was visualised using MEGA 11 (Tamura, et al. 2021). Amino acid sequence motifs and conservation were analysed and visualised using CLC.

### Structure and glycosylation predictions

Structural predictions of A5 were performed using Alpha Fold 3 (Abramson, et al. 2024) using its webserver (https://alphafoldserver.com/). Structures were predicted using the primary amino acid sequence of each protein along with one Zn^2+^ ion, with (Figure 2) or without (Supplementary Figures 1 & 2) a 15-mer ssDNA sequence (5’-GATTACGCACAAACG-3’) derived from the *E. coli rpoB* gene covering the C1576 nucleotide position. Structures were visualised with Mol*Viewer (https://molstar.org/viewer/). Intrinsically disordered protein domains were predicted using the A5 amino acid sequences as queries through the AIUPred server (Erdős and Dosztányi 2024) (https://iupred.elte.hu/) using the default parameters. Glycosylation motifs in each A5 amino acid sequence were predicted using GlycoEP (Chauhan, et al. 2013) (https://webs.iiitd.edu.in/raghava/glycoep/org_index.html). For N-linked glycosylation sites, the binary profile pattern (BPP) approach was used. For O-linked glycosylation sites, the position specific scoring matrix profile pattern (PPP) was used.

### Tissue expression analysis

To assess the relative expression of A5 across different tissues in a marsupial, we analysed A5 mRNA expression in a tissue transcriptome (pooled adult male and female) of the gray short-tailed opossum (GenBank Bioproject: PRJNA200320). To measure mRNA expression, we counted the reads mapping to A5 from the sequence read archive (SRA) of each tissue (Supplementary Table 1) by SRA BLAST (https://blast.ncbi.nlm.nih.gov/Blast.cgi?PAGE=MegaBlast&PROGRAM=blastn&PAGE_TYPE=BlastSearch&BLAST_SPEC=SRA). The query was the opossum A5 coding sequence (GenBank: XM_007501374.1), with the program optimized for highly similar sequences (megablast), and parameters set to: Max targets = 5,000, expect threshold = 1x10^-42^, and other parameters set to their default. The number of reads counted for each sample was normalised across tissues by total library size and the length of the A5 transcript. The A5 expression level in each sample was then normalised against expression of the housekeeping gene RPS13 (GenBank: XM_001366831.1) (de Jonge, et al. 2007), which was counted and adjusted for library size in the same manner as A5. Normalised expression values are the ratio of A5/RPS13 expressed in reads per kilobase of transcript per million mapped reads (RPKM).

### Mammalian and bacterial A5/A3G expression plasmids

The opossum A5, Tasmanian devil A5, and human hA3G proteins used in this study were haemagglutinin (HA)-tagged at their C-terminus and cloned into the mammalian pcDNA3.1 or bacterial pET28a plasmid expression vectors. A5 sequences in pcDNA3.1 were human codon-optimised while in pET28a they were not codon-optimized. The human hA3G plasmid (phu-A3GHA) for expression in mammalian cells has been previously described (Mariani, et al. 2003). Opossum A5 and Tasmanian devil A5 plasmids for expression in mammalian cells (pcDNA3-oA5-HA-HCO and pcDNA3-tdA5-HA-HCO) were codon-optimised and synthesised by GenScript (Singapore). Opossum A5 (pET28a-oA5-HA-OC), Tasmanian devil A5 (pET28a-tdA5-HA-OC), and human hA3G (pET28a-A3G-HA-OC) plasmids for expression in bacterial cells, were synthesised by GenScript. The Vif-deficient HIV-1 virus (HIVΔVif) was expressed from pIIIBDvif (Simon, et al. 1995); Vif-competent HIV-1 (HIV NL4.3) was expressed from pDRNL (Adachi, et al. 1986); and MLV was expressed from pNCS (Bacharach, et al. 2000). The glyco-Gag deficient MLV mutant (MLVΔgg) was generated by mutation of pNCS, to produce pNCSΔgg. A stop codon mutation was introduced into pNCS, in the MLV gPr80Gag reading frame (TAT to TAG) at nucleotide position 608, 12 nt upstream of the normal *gag* ATG, as described previously (Stavrou, et al. 2013). Mutation in pNCSΔgg was verified by Oxford Nanopore plasmid sequencing (Micromon Genomics, Clayton, Australia).

### Mammalian cell cultures

Murine fibroblast (NIH/3T3) cells, human embryonic kidney cells (HEK293T), and human epithelial cervical adenocarcinoma cells modified to constitutively express HIV-1 receptors and coreceptors (TZM-bl cells), were used in this study. TZM-bl cells were obtained from NIH AIDS Research and Reference Reagent Program (Wei, et al. 2002), NIH/3T3 cells were obtained from the American Tissue-type Culture Collection (ATCC, cat# CRL-1658), and HEK293T cells were provided by Richard Axel and verified by the Australian Genomic Research Facility (AGRF, Melbourne, Australia). Cell cultures were incubated at 37°C with 5% CO_2_ using DMEM-10: Dulbecco’s modified Eagle’s medium (Thermo Fisher Scientific, Waltham, MA), supplemented with 10% heat-inactivated fetal bovine serum (Invitrogen, Waltham, MA), 0.02 mM glutamine (Invitrogen), and 1 unit/ml streptomycin (Invitrogen) and 1 unit/ml penicillin (Invitrogen).

### Subcellular localisation

To determine the subcellular localisation of marsupial A5 proteins in comparison to hA3G, HEK293T cells were transfected with 150 ng of pcDNA3-oA5-HA-HCO, pcDNA3-tdA5-HA-HCO, phu-A3GHA, or the empty plasmid expression vector pcDNA3.1 as a negative control. Cells were seeded onto a µ-Slide 8 Well chambered coverslip (ibidi, Fitchberg, USA) at 5.7 x 10^4^ cells per well and grown to approximately 60% confluency at 37°C, 5% CO_2_. Transfections were then performed using Lipofectamine 2000 (Thermo) according to the manufacturer’s protocol, and cells were incubated for 48 hr. Cells were fixed with 4% formaldehyde in phosphate buffered saline (PBS; Thermo) and permeabilized with 0.2% Triton X-100 in PBS.

A5/A3 expression was detected using the following protocol: Fixed cells were first blocked with 3% bovine serum albumin (BSA) for 1 hr. The primary antibody, anti-HA-tag (C29F4) rabbit monoclonal IgG (Cell Signalling Technologies, Danvers, MA), was diluted 1:100 in PBS containing 0.2% Triton X-100 and 3% BSA, and applied to the cells for overnight incubation. Following this, the secondary antibody, anti-rabbit Alexa Fluor 488 (Invitrogen), was diluted 1:500 in PBS containing 0.2% Triton X-100 and 3% BSA, and incubated for 1 hr. Cell nuclei were then stained with Hoechst-33342 diluted 1:5,000 in PBS. All incubations were conducted at room temperature. All stained cells were acquired using a Zeiss Axio Observer fluorescence microscope (Carl Zeiss Microscopy GmbH, Jena, Germany), with a 63x1.4 NA oil immersion lens.

### Rifampicin mutagenesis analyses

Bacterial A3, A5, and negative control expression vectors were transformed into chemically competent C41-pLysS (DE3) *E. coli* and grown on Luria-Bertani (LB) plates with 50 μg/ml kanamycin and 34 μg/ml chloramphenicol (LB-Kan-Cam). Untransformed C41-pLysS *E. coli* were grown on LB plates with 34 μg/ml chloramphenicol (LB-Cam). Individual colonies were selected and grown in LB-Kan-Cam or LB-Cam broth with 1 mM IPTG for 18 hr to reach the stationary phase. Transformed cultures were spread onto LB plates containing 100 μg/ml rifampicin, 50 μg/ml kanamycin, and 34 μg/ml chloramphenicol (LB-Rif-Kan-Cam) or serially diluted and spread onto LB-Kan-Cam plates. Untransformed cultures were spread onto LB-Cam and LB-Rif-Cam plates. Plates were incubated for 18 hr and then mutation frequency was calculated as the ratio of the number of rifampicin-resistant colonies on the LB-Rif-Kan-Cam and LB-Rif-Cam plates to the number on the LB-Kan-Cam and LB-Cam plates.

To identify the specific mutations in the *rpoB* gene of rifampicin-resistant *E. coli*, a 627 nt amplicon was generated by polymerase chain reaction (PCR) of DNA from individual colonies using forward 5’-TTGGCGAAATGGCGGAAAACC-3’ and reverse 5’-CACCGACGGATACCACCTGCTG-3’ primers. Amplicons were purified using the Exo-CIP Rapid PCR Cleanup Kit (New England Biolabs [NEB], Ipswich, MA) and bidirectionally Sanger sequenced (Micromon Genomics) using both amplification primers.

### A5 retroviral restriction analysis

We assessed the antiviral activity of marsupial A5 by examining its capacity to restrict the infectivity of retroviruses, MLV, MLVΔgg, HIVΔVif, and HIV NL4.3. To generate viral particles, HEK293T cells were seeded into 12-well plates at 2.4x10^5^ cells/well. At ∼70% confluency, cells were co-transfected with 5, 50 and 500 ng of oA5 or tdA5 (plus empty vector pcDNA3.1 to a total of 500 ng), or 500 ng of hA3G, and either 500 ng of the MLV expression plasmid pNCS, the MLVΔgg plasmid pNCSΔgg, the HIVΔVif plasmid pIIIBDvif, or the HIV NL4.3 plasmid pDRNL. Cells transfected with only pNCS, pNCSΔgg, pIIIBDvif, pDRNL, or pcDNA3.1 (empty vector) were used as controls. Transfected cells were incubated at 37°C, 5% CO_2_ for 48 hr. Cell culture supernatants containing viral particles were collected and clarified by centrifugation at 1,000 x *g* for 5 min at 4°C. Supernatants were normalised for viral particle content by product-enhanced reverse transcriptase (PERT) assay, as described previously (Vermeire, et al. 2012).

TZM-bl and NIH/3T3 cells were used for infections with HIVΔVif/HIV NL4.3 and MLV/MLVΔgg, respectively. TZM-bl cells were seeded in 96-well plates, NIH/3T3 cells were seeded in 12-well plates. At ∼50% confluency, cells were inoculated with RT-normalised supernatants diluted 1:10 in DMEM-10. TZM-bl cell cultures were incubated at 37°C with 5% CO_2_ and incubated for 48 hr. NIH/3T3 cell cultures were incubated overnight at 37°C with 5% CO_2_, then the viral inocula were removed, cells washed with PBS and fresh media was added. Incubation was then continued for an additional 48 hr. HIV-1 infection of TZM-bl cells was measured by quantification of luciferase activity in cell lysates as previously described (Tyssen, et al. 2010). MLV infection of NIH/3T3 cells was measured by virion-associated RT activity in the cell culture supernatants using the PERT assay as previously described (Hayward, et al. 2020).

### Western blot and deglycosylation analyses

Expression of HA-tagged A5 protein in mammalian and bacterial cells, and the presence of A5 protein in viral pellets, was determined by SDS-PAGE and Western blot. Mammalian cell lysates and viral pellets were collected from infected cell cultures as previously described (Hayward, et al. 2022). Bacterial cultures were generated as described above in ‘Rifampicin mutagenesis analyses’. After reaching the stationary phase, 5 ml of each culture was pelleted and lysed with 200 μl of Triton-Tris lysis buffer containing 0.1% Triton X-100 (Sigma-Aldrich, Burlington, MA) and 10 μg/mL each of aprotinin, leupeptin, and pepstatin A protease inhibitors (Sigma-Aldrich).

Proteins in all lysates were analysed by SDS-PAGE under reducing conditions with 10% beta-mercaptoethanol (Sigma-Aldrich). To deglycosylate proteins, samples were treated with PNGase F (NEB) following the manufacturer’s instructions. Proteins were transferred to a nitrocellulose membrane (Sigma-Aldrich). For the Western blot analysis, the primary antibodies were (i) a rabbit anti-HA monoclonal antibody (C29F4, NEB), (ii) a mouse anti-HIV-p24 monoclonal antibody (NIH-183, NIH HIV reagent program), and (iii) a rabbit anti-MLV-Gag polyclonal antibody (ab100970, abcam), all diluted at 1:1,000 with 0.1% Tween-20, 5% BSA, 0.02% azide in Tris-buffered saline (TBS) and incubated overnight at 4°C. The secondary antibodies were (i) a goat anti-rabbit IRDye 800CW fluorophore (LI-COR, Lincoln, NE) and (ii) a goat anti-mouse Alexa Fluor 680 fluorophore (Invitrogen), both diluted at 1:10,000 with 0.1% Tween-20, 5% BSA, 0.02% azide in TBS, incubated at room temperature for 1 hr. For visualisation, membranes were scanned with a ChemiDoc MP Imaging System (Bio-Rad, Hercules, CA), at 600 and 800 nm.

### Hypermutation and molecular clock analyses

The presence of A3-like cytosine deamination on ancient endogenised opossum retroviruses was assessed by hypermutation analysis (Lee, et al. 2008) as previously described (Hayward, et al. 2018). ERV sequences were extracted from the *Monodelphis domestica* genome assembly, mMonDom1.pri (GenBank: GCF_027887165.1). Full-length endogenous betaretroviruses and gammaretroviruses were identified in the opossum genome as described previously (Hayward, et al. 2013), using the Jaagsiekte sheep retrovirus (JSRV, GenBank: NC_001494) and koala retrovirus (KoRV-A, GenBank: AF151794) sequences, respectively. Unique target site duplications were identified for each ERV, confirming that each was the product of a separate retroviral integration event, rather than a chromosomal duplication event (Supplementary Table 2). Estimations of the time since integration for each ERV were calculated by molecular clock analysis as previously described (Hayward, et al. 2013) using a mutation rate of 0.30x10^-8^ mutations per nucleotide per year for *Monodelphis domestica* (Peña-Garcia, et al. 2024).

Multiple sequence alignments (MSAs) of ERVs were generated using MUSCLE (Edgar 2004). Phylogenetic analyses of ERVs were undertaken with the maximum likelihood method and the best-fit model (GTR+G+T) with 1,000 bootstrap replicates in CLC. A sub-clade of closely related ERVs, and more distantly related outgroup ERV, was selected for each group for further analysis. The ancestral sequence of each group of ERVs was computationally extrapolated using MEGA 11 (Tamura, et al. 2021) using the accumulated mutations in each ERV through estimation of the ancestral state of each node in the tree. ERVs were then aligned against the ancestral sequence using MUSCLE (Edgar 2004) and gaps relative to the ancestral sequence were removed from the alignment. The ERVs and extrapolated ancestral sequences are provided in Supplementary Data 1.

A3-like mutational biases were evaluated by analysis of the alignment of each group of ERVs with its extrapolated ancestral sequence using the Hypermut2 tool (https://www.hiv.lanl.gov/content/sequence/HYPERMUT/hypermutv2.html) (Rose and Korber 2000) with context enforced on the ancestral reference sequence. Fisher’s exact test was used within the Hypermut2 tool to determine statistical significance.

## Acknowledgements

We thank Elsie Williams for providing the C41-pLysS *E. coli,* Richard Axel for providing HEK293T cells, Stephen Goff for providing pNCS, and Tom Hope for providing phu-A3GHA and pIIIBDvif. J.A.H, S.T, P.E, and this study were funded through the kind support of the Estate of GWA Griffiths. J.A.H and this study were additionally funded by the Upotipotpon Foundation, Miss Gwen Hotton, Mrs Rosemary Jacoby, and the Jim and Margaret Beever Fellowship. We gratefully acknowledge the contribution of the Victorian Operational Infrastructure Support Program, received by the Burnet Institute.

## Data Availability

ERVs and extrapolated ancestral sequences used in the hypermutation assay are included in Supplementary Data 1.

